# Freshwater connectivity transforms spatially integrated signals of biodiversity

**DOI:** 10.1101/2022.07.20.500822

**Authors:** Joanne E. Littlefair, José S. Hleap, Vince Palace, Michael D. Rennie, Michael J. Paterson, Melania E. Cristescu

## Abstract

Aquatic ecosystems offer a continuum of water flow from headwater streams to inland lakes and coastal marine systems. This spatial connectivity influences the structure, function and dynamics of aquatic communities, which are among the most threatened and degraded on earth. Environmental DNA achieves biodiversity surveys in these habitats in a high-throughput, spatially integrated way. Here, we determine the spatial resolution of eDNA in dendritic freshwater networks that are typical of aquatic habitats. Our intensive sampling campaign comprised over 430 eDNA samples across 21 connected lakes, allowing us to analyse detections at a variety of scales, from different habitats within a lake to entire lake networks. We found strong signals of within-lake variation in eDNA distribution reflective of typical habitat use by both fish and zooplankton. Most importantly, we also found that connecting channels between lakes resulted in an accumulation of downstream eDNA detections in lakes with a higher number of inflows, and as networks increased in length. These findings have profound implications for the interpretation of eDNA detections in aquatic ecosystems in global-scale biodiversity monitoring observations.

## Introduction

In freshwater, the direction and strength of water flow among habitats shapes processes of recolonization, genetic diversity, adaptation, ecological flows, and facilitates population resilience^1,2^. Within lakes or rivers, there is further partitioning according to fine-scale environmental variables such as water temperature and oxygen concentration; some organisms are adapted for life in the littoral zones of lakes while others specialise in the deeper waters of the pelagic zone. Given its importance, spatial connectivity has become a major concern in conservation when designing habitat management plans for threatened freshwater populations which are declining at a catastrophic rate^3–5^. Historically, genetic data obtained directly from inhabiting organisms has provided valuable information for inferring the biotic connectivity of freshwater habitats^6^. The non-destructive and non-invasive nature of environmental DNA (eDNA) sampling is important in contributing to species conservation goals and cultural sensitivities, as many communities reject traditional lethal survey netting^7^. However, the wider adoption of species detection based on eDNA by researchers, managers and policy makers depends heavily on our ability to accurately interpret eDNA signals, particularly when trying to distinguish organisms that currently inhabit a particular habitat, from those that inhabit nearby habitats, or organisms previously but no longer occupying the area^8^. This is particularly relevant for conservation and biodiversity projects, for example, when assessing the presence of rare or invasive species.

It is generally accepted that the complex “natural history” of environmental nucleic acids combined with prevailing environmental and hydrological conditions can influence the spatial or temporal resolution of species detection. Spatial scales of detection are likely to be an interaction between the rate of initial local production of eDNA, combined with subsequent dilution and transport in the environment (including vertical settling), and eventual degradation of DNA molecules^9,10^.

Situations with weak transport effects (perhaps combined with high dilution) will produce local signals that rapidly dissipate in strength further from the source population. In these cases, eDNA has shown high spatial fidelity with visual or trap-based surveys^11^. For example, harbour porpoise eDNA could not be detected further than 10m away from the animals due to dilution effects^12^. In contrast, when prevailing environmental conditions produce strong transport effects, possibly combined with high rates of initial eDNA production or low rates of dilution, eDNA signals can be transported away from the initial source population of animals, signalling regional, rather than local, biodiversity^8^. To date, studies investigating eDNA transport have concentrated on downstream movement in lotic systems, which varies with the velocity of river flow but is likely to be a significant force in shaping nucleic acid distribution^13–15^. Other types of water movement, such as hydrological forces within lakes have been less well studied (although see^16,17^).

Thus, the accurate spatial interpretation of eDNA-based surveys in aquatic networks depends on explicitly modeling the retention and flow dynamics of eDNA away from local habitats on a landscape scale.

Here, we used an eDNA metabarcoding approach to analyse zooplankton and fish communities in three lake networks containing 21 Canadian boreal lakes connected only by surface flow, quantifying the spatial distribution of eDNA signals within and among lakes, as well as among networks. We validated eDNA-based results against both current and historical population records collected since the 1970s. To investigate within-lake patchiness in eDNA signals, we examined how eDNA sampled in different zones of the lake matched known habitat use by animals (i.e., littoral, epilimnetic and deep-water). We also evaluated the frequency of downstream detection of eDNA and the influence of hydrological factors. To investigate between-lake variation in eDNA signals, we classified eDNA based on whether it matched historical and current population records as either expected or unexpected. Finally, we propose a series of spatially-explicit models in which the arrangement of lakes within a network influences the dynamics of eDNA, using a patch dynamics perspective^18^. In all these systems, we assumed that each lake had the same local production of eDNA. Proposed models of within-network eDNA transport (Figure 1A) are as follows:

**Figure 1A:**
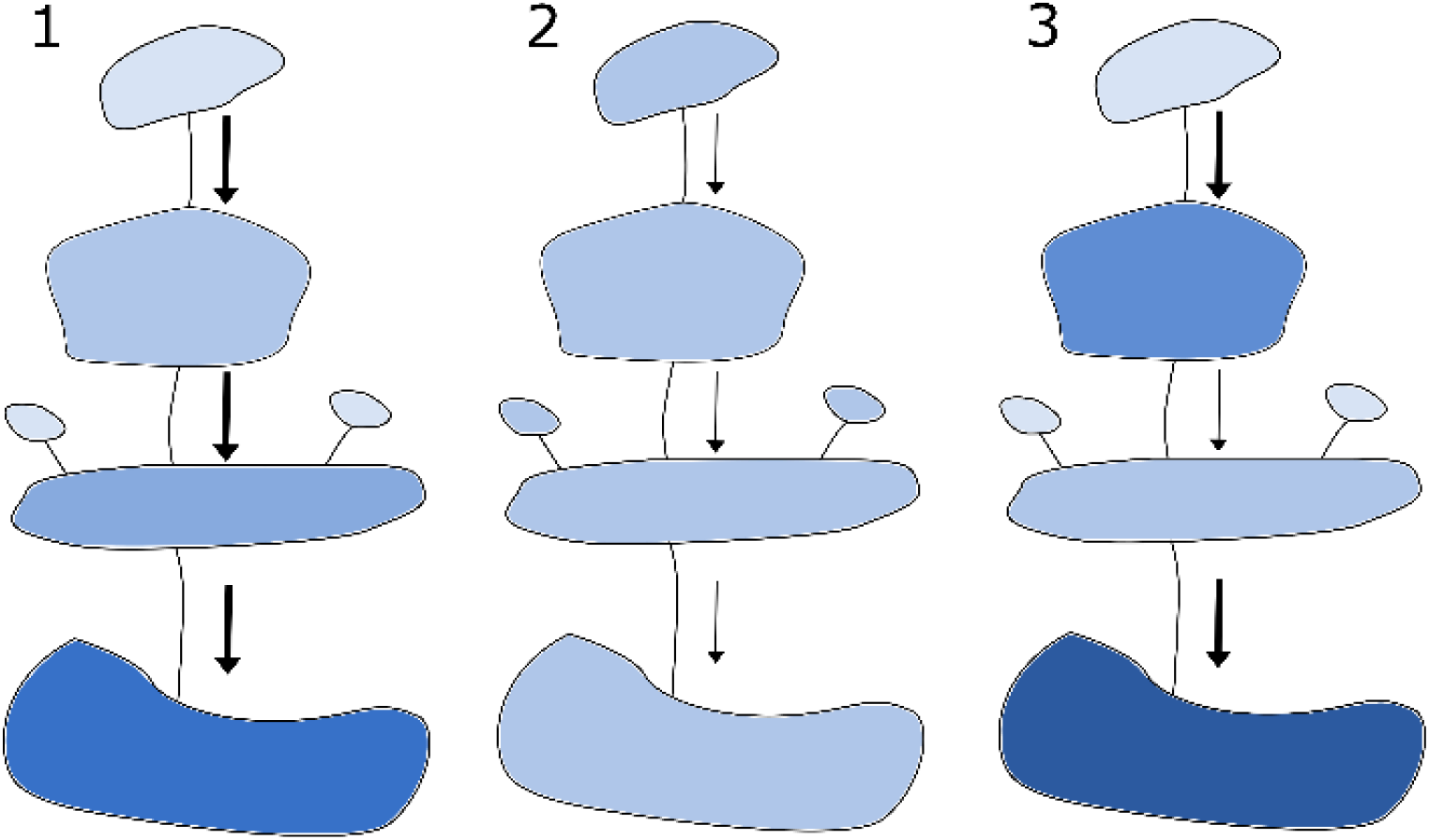
We propose a ‘spatially explicit model’ in which the arrangement and the size of patches (lakes) influences the dynamics of eDNA. We assume a movement of DNA directional to the flow. For simplicity we can assume that all patches can sustain equally diverse communities (as in patch dynamics perspectives^18^).Proposed models are as follows: 1)High flow networks leading to short retention time for eDNA and high water turnover within patches; there is insufficient retention time to ensure the degradation of eDNA within the site. Prediction: eDNA reflects **regional** rather than local diversity, and unexpected detections increase with increasing lake connectivity. 2)Low flow networks leading to higher retention time of eDNA within patches and slower water turnover within patches: most of the eDNA signal will degrade within the resident patch. Prediction: the genetic diversity recovered by eDNA in downstream patches corresponds mainly to **local** diversity rather than regional diversity. 3)Mixed networks in which patches can have different flow regimes, some with low and some with high retention time. The shorter the retention time the more likely the flow of eDNA signal downstream. Prediction: some parts of the network retain **local** signal and more dynamic parts retain **regional** signal. In the figure darker shading represents higher rates of unexpected detections with eDNA and a bolder arrow represents higher water flow between patches.

**Figure 1B:**
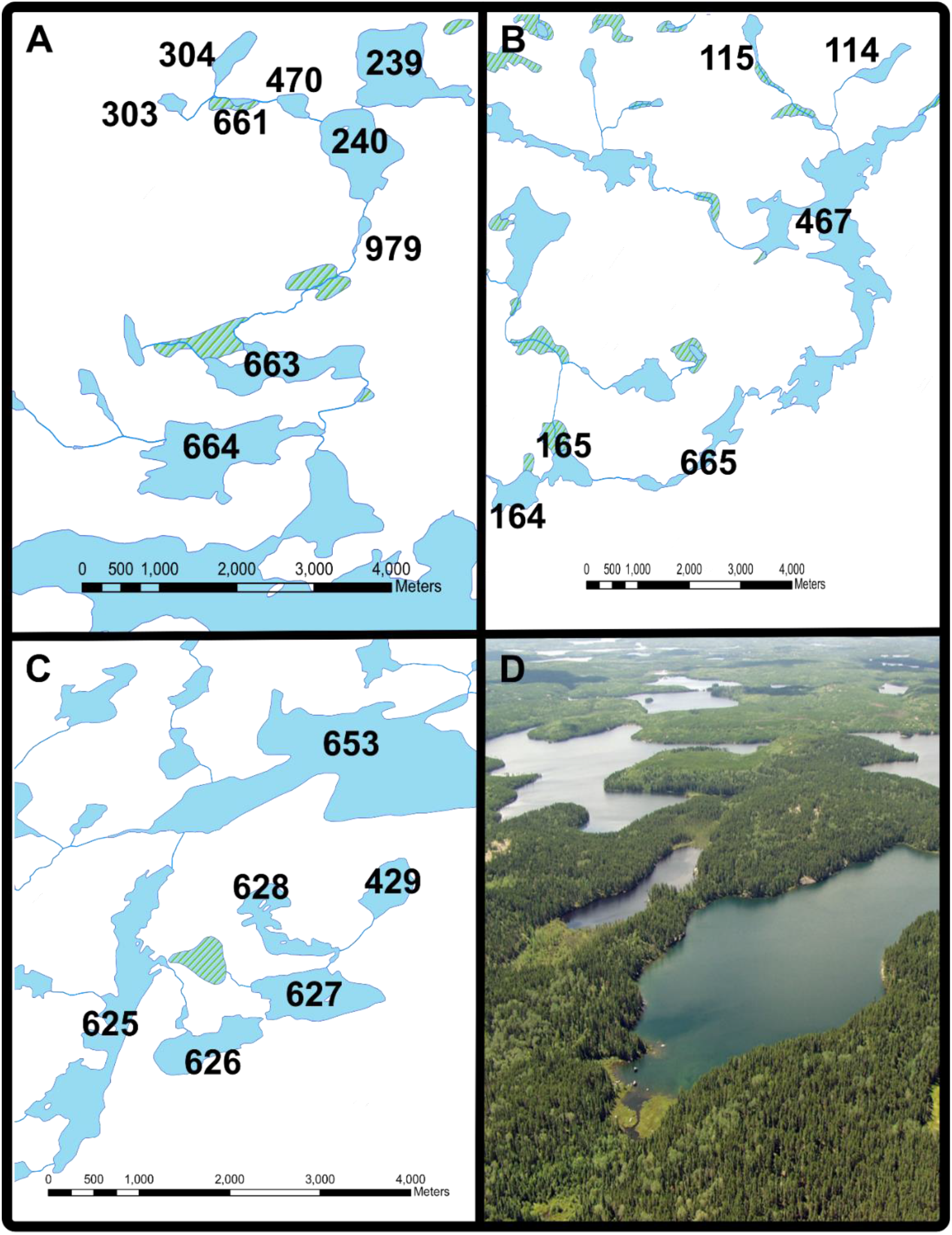
Maps of the three lake networks and aerial view of the connected lake and stream habitats of the Experimental Lakes Area, Ontario (Panel A – Chain 1, Panel B – Chain 2, Panel C – Chain 3). Each lake at the Experimental Lakes Area has a unique identification number, which are represented on the map. An aerial photograph shows the connected lakes of Chain 1 (Panel D).

1)High-flow networks leading to short retention time for eDNA and high water turnover within lakes; there is insufficient retention time for the degradation of eDNA within the site. Prediction: unexpected detections (detection of species that do not reside in the particular habitat) increase gradually with lake chain number and eDNA reflects regional rather than local diversity.

2)Low flow networks with slow water turnover, leading to higher retention time for eDNA within lakes and degradation of most of the eDNA signal within resident lakes. Prediction: eDNA detections correspond to local diversity rather than regional diversity and the rate of unexpected detections remains relatively constant throughout the system (both within and among lakes).

3)Mixed networks in which lakes have varied water retention times. The shorter the retention time, the more likely the eDNA signal will flow downstream. Prediction: some parts of the network retain local signal and more dynamic parts retain regional signal. Unexpected detections vary according to the local flow regime.

## Results

We detected all fish species with eDNA that were recorded by conventional techniques at the IISD Experimental Lakes Area. We also made additional detections of *Esox masquinongy*, which is known to exist regionally. After controlling for sequences detected in blank samples and mock communities, we made 1,909 detections across the dataset, with an average of 4.6 species detections per sample (Extended data table 1). Sample accumulation curves for every lake were reflective of good sampling coverage using the 12S marker, based on our assessment of the plateaued sampling curves (Extended data figure 1). Of detections found in the lake samples (i.e. shoreline, deep-water and pelagic-surface samples), 67% were validated by conventional current and historical fish monitoring records. Those that were predicted by conventional monitoring methods had significantly higher per-sample ASV abundances (Quasipoisson GLM, p < 0.001, predicted by fishing records median = 949, IQR: 156-3267 Not predicted by fishing records, median: 3, IQR: 1-122).

We made 6630 zooplankton detections with the COI dataset with an average of 31.8 ASVs per sample that could be assigned to class level or below. Sample accumulation curves indicated that sampling coverage was not as extensive as that observed for fish (Extended data figure 1). The COI marker detected many other taxa that are not considered zooplankton and were excluded from further analysis; primarily insects (Extended data figure 2). Of 264 zooplankton ASVs detected, 36 matched to Calanoida, 33 to Cyclopoida, 67 to Cladocera, 23 to Diptera, and 100 to Rotifera.

### Within-lake variation in eDNA distribution

We detected different fish community compositions among different habitats within the lakes (e.g. shoreline, deep-water, pelagic-surface transect, in- and out-flows, PERMANOVA R^2^ = 3.69%, p < 0.001). Hypothesis testing showed that the interaction between fish habitat preference and eDNA sample location was a significant predictor of eDNA ASV abundances (X = 85.8, p < 0.001). In general, ASVs were most abundant when eDNA sample location matched known fish habitat preferences. In particular, ASV abundances from profundal species were highest in deep-water samples; these species were infrequently detected by shoreline or pelagic samples when these species were known to be present in the lake (Figure 2A, Extended data figure 3). Littoral-benthic ASVs were much more abundant in the shoreline samples, and a greater diversity of littoral species were found with the shoreline samples than any other sample type in the largest lakes (Extended data figure 3). Generally, the largest, stratified lakes in our study had the greatest distinctions between community compositions found in different sample types (Extended data figure 3).

**Figure 2:**
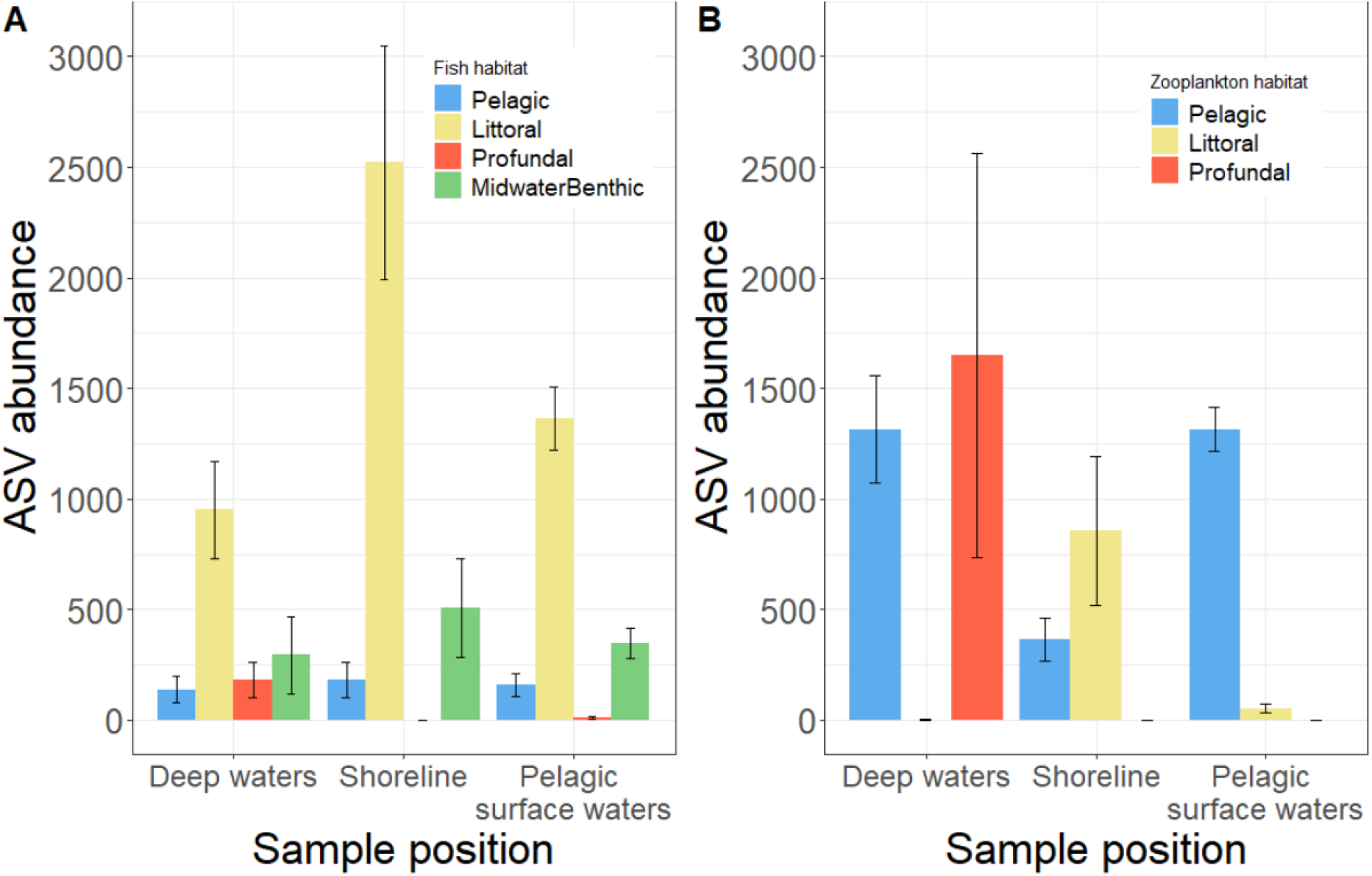
Fish eDNA ASV count is influenced by the interaction between sample location (deep water, shoreline or pelagic-surface transect) and classification of fish habitat (littoral-benthic, midwater-benthic, profundal or pelagic). eDNA ASV counts reflect the fish species using those habitats. Fish with a profundal habitat preference were principally found in deep water samples while littoral-benthic fish were predominantly detected in the shoreline samples. Zooplankton eDNA read count was also influenced by the interaction between the location at which the samples were collected and the habitat classification of the zooplankton. eDNA ASV counts reflects the zooplankton species’ habitat use. Zooplankton species with a profundal habitat preference were principally found in deep water samples while littoral zooplankton was predominantly detected in the shoreline samples.

We also found strong spatial structure in zooplankton ASV abundance. The interaction between eDNA sample location and zooplankton habitat use was a significant predictor of ASV abundance (Χ = 168.6, *p* < 0.001). Deep-water samples had larger counts of hypolimnetic zooplankton, but no littoral species. Hypolimnetic zooplankton ASVs were not abundant in the pelagic-surface or shoreline samples. Samples taken at the shoreline were best at detecting littoral zooplankton, which were rarely found in other locations, and samples in the pelagic-surface transect were best at detecting pelagic species (Figure 2B, Extended data figure 4).

### Between-lake variation

Modelling demonstrated increased numbers of unexpected eDNA detections in lakes as connectivity increased. Here we included lake chain number as a proxy for connectivity, but the same results applied to connectivity described as an increasing number of inflows (see methods). For the fish dataset, there was a significant interaction between count type (i.e. expected or unexpected eDNA detections, as defined by detection compared with conventional surveys) and the position in the lake chain (Figures 3 and 4, GLMM, LRT = 12.56, *p* = 0.0004) on the number of detections per sample. While unexpected detections moderately increased in lakes further down the networks, expected detections remained roughly constant throughout the networks (Figure 4). These unexpected detections mostly matched species living in the lake directly upstream to the one being sampled (Figure 3). In the zooplankton dataset, the interaction between count type and lake chain number was not significant. Instead, there was a significant main effect of lake chain number on all detections, with both expected and unexpected detections increasing downstream (Figures 3 and 4, GLMM, LRT = 4.88, *p* = 0.027).

**Figure 3:**
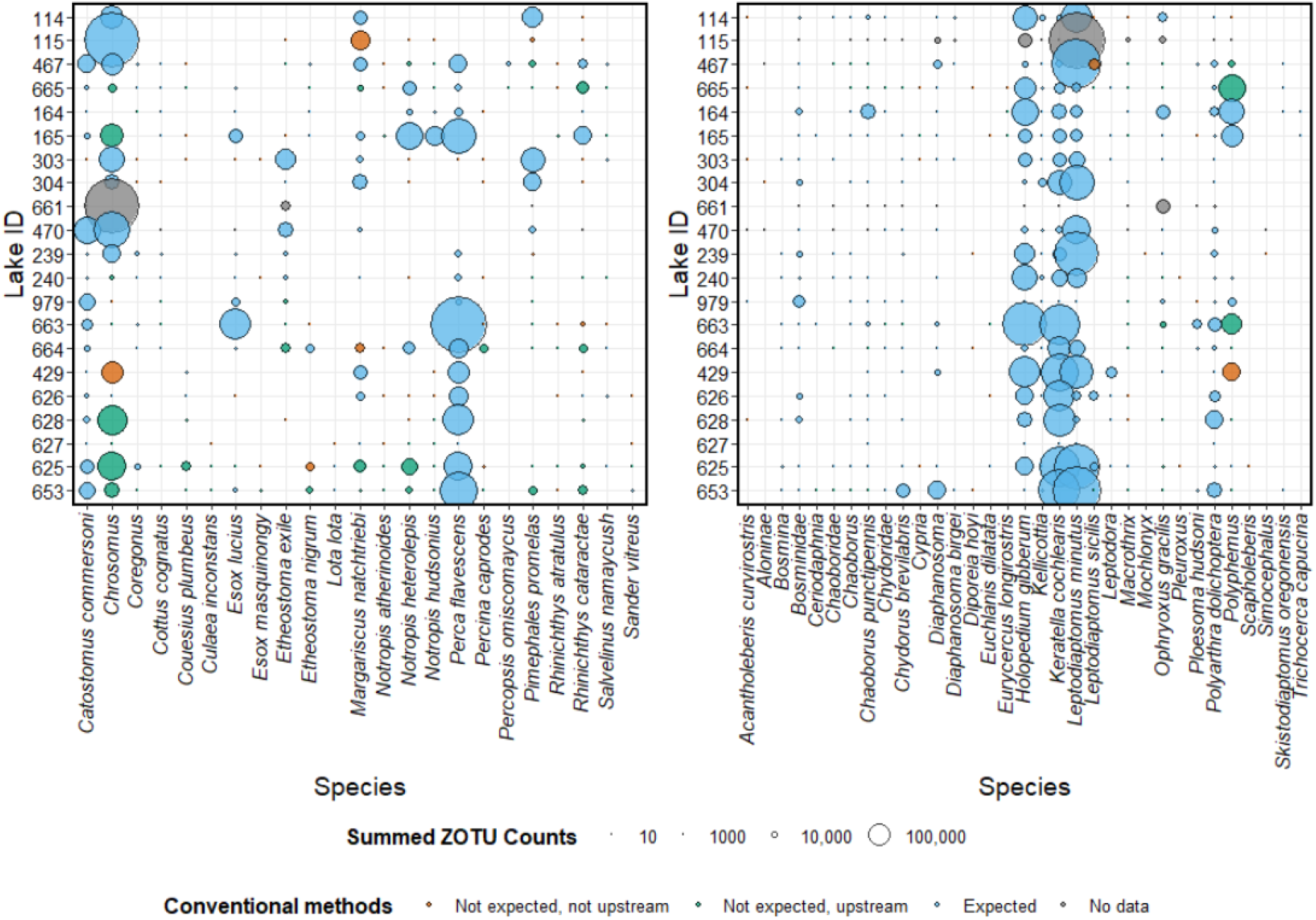
Bubble plot to show eDNA detections of each species in each lake. Lakes are ordered by lake network, and within each lake network they are ordered by their position within the network beginning with headwater lakes. Bubbles represent the summed ASV counts for all the samples associated with that lake, including the inflows and outflows. Bubble size is weighted by the number of ASV counts. The colour of the bubble compares eDNA detections to detections with conventional monitoring methods. Blue = expected according to conventional monitoring, green = not expected according to conventional monitoring but present in the lake immediately upstream (according to either monitoring method), orange = unexpected according to conventional methods, grey = no data on whether species is expected or unexpected because conventional monitoring is not adequate in that habitat.

**Figure 4:**
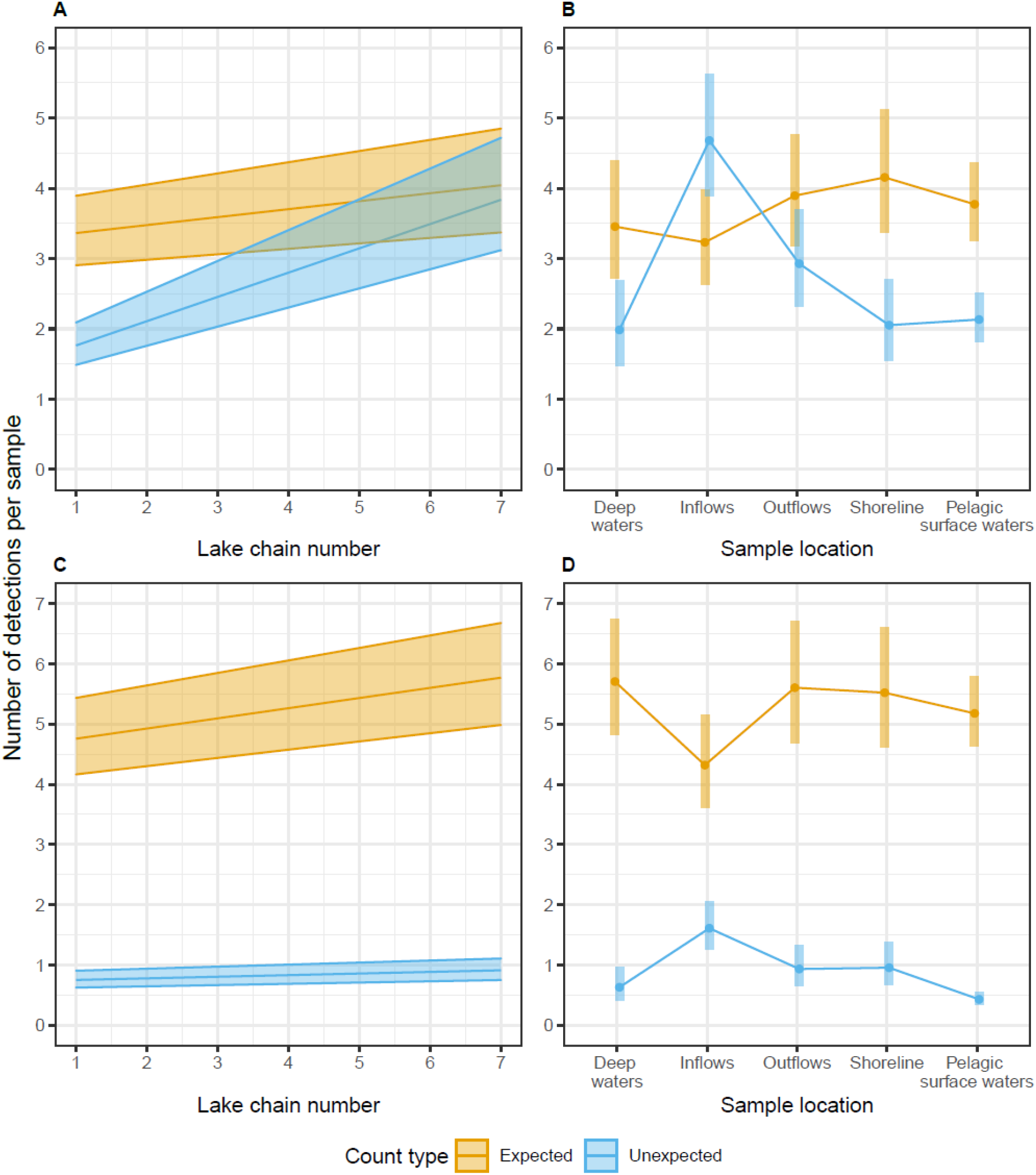
Number of per-sample expected and unexpected eDNA detections as a function of lake chain number and sample location for fish (A, B) and zooplankton (C, D) datasets. Unexpected eDNA detections are those not matched by historical and current fishing and zooplankton records. Across all three networks, there is a pattern of an increase in unexpected fish species detections in downstream lakes, which is not reflected by expected detections. Plots show back-transformed model predictions from negative binomial GLMMs built in glmmTMB.

There was a significant interaction between sample location and count type on the number of species detected, meaning that both expected and unexpected detections were not distributed evenly throughout the freshwater networks for either fish or zooplankton eDNA (fish GLMM: LRT = 42.7, *p* < 0.001; zooplankton GLMM: LRT = 79.8, *p* < 0.001). When considering fish eDNA detections that are expected when compared to conventional methods, all sample locations detected equal species richness (Figure 4). This was also broadly similar to the zooplankton eDNA detections, except that there was a difference between the zooplankton deep water and inflow locations in the post-hoc testing, with inflows detecting fewer species matching the conventional survey (5.5 taxa in deep water samples versus 4.1 in inflow samples, *p* = 0.024). When considering fish eDNA detections not matched by conventional methods, post-hoc tests showed a higher number of these types of detections in inflows (mean 4.8 species) when compared with shoreline (2.1 species, *p* < 0.001), deep water (2.0 species, *p* < 0.001) and pelagic-surface (2.2 species, *p* < 0.001) samples (Figure 4). Zooplankton eDNA behaved similarly, with greater amounts of these mismatched detections in certain areas of the lake (GLMM, LRT = 79.8, *p* < 0.001). Specifically, higher numbers of mismatched detections were detected in the inflows (1.5 taxa) compared to the deep water (0.61 taxa, *p* = 0.011), the outflows (0.85 taxa, *p* = 0.023), the shoreline (0.87 taxa, *p* = 0.047), and the pelagic-surface (0.40 taxa, *p* < 0.001). There were also significant differences between the outflows and pelagic-surface samples (*p* = 0.002), and between the shoreline and pelagic-surface samples (*p* = 0.002).

### Stream discharge and eDNA detections

Inflow discharge did not interact with count type to influence the numbers of fish eDNA detections found in the inflow samples (GLMM, LRT = 0.0001, *p* = 0.99). Moreover, there were no significant main effects of inflow discharge (GLMM, LRT = 0.0003, *p* = 0.996) or count type (GLMM, LRT = 1.06, *p* = 0.304) on the number of eDNA detections. The zooplankton dataset displayed a different pattern, with a significant interaction between inflow discharge and count type (GLMM, LRT = 4.02, *p* = 0.045); as the inflows increased in discharge, the number of expected counts per sample increased. Unexpected counts were lower than expected counts overall, and slightly decreased as stream discharge increased.

## Discussion

Our intensive sampling campaign involving 430 samples across 21 connected lakes allowed us to analyse eDNA detections at multiple scales, spanning different habitats within a lake to entire lake networks. Our results emphasise the importance of considering aquatic connectivity – both within lakes and flowing connections between lakes – in shaping the distribution of molecular detections. Given that 9.5% of known species occupy freshwater habitats, including one third of the world’s vertebrates, and that freshwater habitats are characterised by connectivity, our results have major implications for the application and interpretation of eDNA to characterise, survey and monitor aquatic communities^4^.

Lakes with greater connectivity (i.e., those further downstream or with a greater number of inflows) had more unexpected detections of fish eDNA. This analysis only included data from samples collected within the lakes themselves (i.e., not samples collected in inflow and outflow streams or surrounding ponds), thus precluding the contribution of a greater number of inflow samples to the accumulation of additional detections. Occasionally, the unexpected eDNA signal was very strong, being found in the majority of samples from a lake. This suggests `that eDNA is detecting animals that have not been caught with conventional techniques. Despite these few occurrences, long-term ecological research sites are some of the best places in the world to perform this kind of comparison between molecular and conventional fishing techniques, because of the comprehensive sampling effort and extensive species databases covering many lakes. It is likely, therefore, that eDNA from upstream lakes is transported via the inflow and then becomes mixed with the downstream epilimnion. While we might expect downstream lakes to have a greater species richness generally due to increasing size, downstream lakes were not consistently the largest in our study. Moreover, the rate of increase in unexpected detections was much higher than the moderate increase in expected detections further down the networks. Previously, invertebrate eDNA has been shown to travel between 9 and 12km downstream from a river flowing downstream from a lake^13^. Other studies have also shown the transport and accumulation of eDNA on the scale of several kilometres^8,19–22^. With regard to our model scenarios, this points to a model similar to scenario 1 – the propensity for the downstream accumulation of DNA, in which lower retention times do not allow for the complete degradation of eDNA within a lake and thus molecular signal accumulates with greater freshwater connectivity.

Inflows and outflows create fine-scale physical, chemical and biological heterogeneity across lakes^23^. This also applies to eDNA, as we found heterogeneous eDNA signals in both the inflows and outflows of lakes. Sometimes small fish and littoral zooplankton live in streams, but we also found eDNA originating from other fish species that would not normally dwell in streams. As a result of these two phenomena, species richness detected in inflows was high compared with samples from other locations. Moreover, there were large amounts of “unexpected” detections in inflow samples that did not match the composition of the receiving lake. In some instances, it was clear that non-resident eDNA was flowing from the inflow and creating a plume of DNA into the downstream lake. For example, the inflow of lake 665 contained large amounts of DNA from pearl dace (*Margariscus natchtriebi*) and longnose dace (*Rhinichthys cataractae*), both of which are residents of upstream lake 467 (Chain 2, Figure 1B). Likewise, the inflow of lake 979 contained eDNA from non-resident Iowa darter (*Etheostoma exile*), which is a resident of upstream lake 240. The pelagic-surface sampling points closest to these inflows in lake 665 and lake 979 have a similar community composition to the water sampled at the inflow, but this non-resident signal is not apparent in further pelagic-surface sampling points. This plume of non-resident eDNA might be facilitated by the fact that both receiving lakes are reasonably narrow “channel”-shaped lakes coupled with a proportionally large inflow. It is possible that morphometry combined with the degree of connectivity is an understudied determinant of the level of incursion from residual molecular signal.

Although the streams at IISD-ELA are relatively small, they are also short in length, and can therefore contribute to downstream eDNA signals before the molecules degrade. The complexity of interacting factors may explain the lack of a simple relationship between discharge and eDNA detections that is consistent across fish and zooplankton eDNA. Intuitively, we might expect increased stream discharge to carry eDNA further downstream^24^. However, increased water volume has also been shown to have a moderately diluting effect on eDNA copy number^25,26^. Flow regime might also act indirectly on eDNA detectability by affecting other abiotic factors, such as the levels of inhibitors in the water or degree of particle settling or resuspension from the streambed, which influences the amount of eDNA detected^15,27^. Other studies hint at a seasonal effect on eDNA transport in inflows and accumulation in downstream habitats; at the time of sampling, some inflows were slow flowing as it was the height of summer. While some flash rain events did occur during our six-week sampling period, stream flow is typically highest during spring snowmelt in this region. However, simply measuring stream discharge at single time points may be less likely to reflect the potential transport of eDNA among lake ecosystems, compared to a more integrated picture of ongoing eDNA accumulation in downstream lakes.

We have demonstrated robust evidence for the spatial partitioning of DNA signals within a lake. Largely, the habitat preferences of fish and zooplankton defined the community composition of the eDNA signals found in those sample locations. Strikingly, the thermocline seems to be an important factor in restricting eDNA flow to surface waters, as hypolimnetic species are almost exclusively detected in profundal cold water. During the summer months, some zooplankton e.g., *Leptodiaptomus sicilis* and fish species like lake trout (*Salvelinus namaycush*) and slimy sculpin (*Cottus cognatus*) are isolated to profundal cold water below the thermocline due to their oxythermal habitat requirements, and we found that their eDNA was almost exclusively detected in those respective environments. This is a seasonal pattern driven by stratification of lakes, which isolates both cold water species and their eDNA to the bottom of lakes during the warmer months^16,28,29^, resulting in little hydrologic connectivity between the epilimnion and hypolimnion. Studies using radioactively-labelled water added to similar sized lakes showed that there is very limited diffusive exchange across the thermocline and concluded that the thermocline acts as a barrier inhibiting the downward transfer of turbulent mixing energy^30,31^. This could explain the infrequent detection of DNA belonging to cold-water species in any other parts of the lake. Some of our study lakes were too shallow to stratify. Our sampling design still incorporated a water sample from the bottom of these small, shallow lakes, but in these cases the detected community composition was more similar to samples from the shoreline and pelagic-surface waters (Extended data figures 3 and 4). In large, stratified lakes, however, the detected community composition differed greatly between deep water samples and shoreline/pelagic-surface samples.

eDNA from littoral fish species was spatially structured between the shoreline and pelagic samples, with samples at the shoreline containing more littoral fish sequences such as fathead minnows (*Pimephalas promelas*), yellow perch (*Perca flavescens*) and blacknose shiner (*Notropis heterolepis*). This pattern was even more pronounced with littoral zooplankton, which were very rarely detected in pelagic or deep-water samples. In particular, *Polyphemus pediculus*, a predatory littoral cladoceran, had very high sequence numbers in shoreline samples. We found the separation of distinct littoral and pelagic communities surprising, because radio-tracer experiments in IISD-ELA lakes have shown that the epilimnion is fully mixed within one day of tritiated water injection^31^, due to wind stress on the lake surface. However, the heterogeneous eDNA signal originates from the larger lakes in our study, where distinct community compositions were detected between shoreline and pelagic epilimnion samples, reflecting the habitat preferences of these species (Extended data figure 4). These larger lakes likely support unique littoral and pelagic fish and zooplankton communities, as well as present longer times for eDNA signals to mix across the epilimnion. Other eDNA studies from single lakes have hinted at this finding; for example, small littoral fish were found to have a greater relative sequence abundances in shoreline samples compared to samples from the centre of large lakes (1480ha^29^; 122ha and 4343ha^32^).

Within lakes, heterogeneity in species detection with eDNA is shaped by heterogeneity in habitat and thermal structure. Especially in larger lakes, eDNA signals had spatial structure that reflected the habitat preferences of animals. There is also clear evidence that eDNA can reflect upstream communities of organisms when a high degree of ecosystem connectivity is present. Using a landscape perspective of freshwater ecology, lakes are explicitly viewed as connected to each other and their catchment area. eDNA does not accumulate homogeneously downstream, but both landscape factors (i.e., the position of the lake relative to others in the network) as well as individual lake-specific factors such as morphometry influence the degree of eDNA signal. We have highlighted how motion in water, which is a fundamental process in freshwater systems, will shape detectable eDNA signals and therefore biomonitoring sampling designs.

## Acknowledgements

Many IISD Experimental Lakes Area researchers and students helped to generate the field data for which we are very grateful. S. Michaleski, and R. Henderson contributed specific field assistance to this project. We thank D. Schoen for laboratory space and K. Sandilands for advice regarding lake hydrology. L. Veilleux, N. Vachon, H. Massé, and N. Tessier from the Ministère des Forêts, de la Faune et des Parcs (Québec) contributed the fish tissue samples for the mock community. J. Carreau and P. Lafrance at WSP Montréal provided helpful discussions regarding this project. This work was funded by a Mitacs Accelerate Industrial Fellowship (JEL and MEC), NSERC Engage award (MEC) NSERC Discovery grants (MEC, MDR), Québec Centre for Biodiversity Science Excellence award (JEL), Experimental Lakes Area fee rebate, and the WSP Montréal, Environment Department and a Canada Research Chair award (MDR).

## Data accessibility

All data have been deposited in DataDryad under doi:10.XXX/XXX.XXX The bioinformatic pipeline from adapter removal to sample table creation can be found in the github repository https://github.com/CristescuLab/YAAP, as ASV_pipeline.sh.

## Methods

Sampling for eDNA was conducted from June to July 2017 at the International Institute for Sustainable Development Experimental Lakes Area (IISD-ELA), Ontario, Canada, a facility for whole-lake ecosystem experimentation and monitoring. Situated on the Canadian Shield, a geological formation dominated by granite bedrock, the region is characterized by a high density of lakes linked primarily by surface water flow with negligible groundwater flow. We collected 430 water samples from three lake chains composed of 21 lakes ranging in size from 2-210 hectares (Figure 1B; Chain 1 = 9 lakes; Chain 2 = 6 lakes; Chain 3 = 6 lakes). We selected lake chains with the most complete historical population records of fish and zooplankton communities. Conventional monitoring and enumeration of fish and zooplankton populations in these lakes has taken place with varying levels of intensity since the 1960s. Lakes in each chain were connected by streams of varying flow regimes. We characterised lakes according to lake chain number, which measures landscape position relative to other lakes, linearly connected through surface flow^33^.

Several lakes have been monitored annually to bi-annually for fish (spring and fall sampling) using a combination of non-lethal gillnetting and trapnetting^34,35^ (Extended data table 2). All other lakes have been surveyed 1-2 times using a set of experimental gillnets, trap nets and minnow traps^36^. Since 2014, several lakes have been sampled using a modified version of the Ontario Ministry of Natural Resources Broadscale Monitoring method, which applies both North American and Ontario small mesh nets in an area-weighted fashion across depth strata^37^.

Between 1968 and 2017, many of the study lakes have been sampled for zooplankton using Schindler-Patalas traps^38^, nets, and tube samplers^39^. Several of the study lakes have been sampled biweekly during the open-water season for up to 53 years. On most sampling dates, a minimum of 300 animals were identified to species level. For full details of historical population monitoring, see Supplementary Note 1. We took additional zooplankton hauls in 2017 to ensure that all lakes had current species richness information, using a 30cm diameter net with 53 μm mesh lowered to 1.5m above the lake bed. Samples were preserved in formalin for morphological identification^40^.

### eDNA collection and analysis

Each of the 21 lakes were sampled for eDNA using a variety of strategies. To evaluate spatial differences across lakes, a transect of 500ml pelagic-surface samples was taken at five evenly spaced intervals across the lake, including the deepest point of the lake (n = 5). To evaluate depth-specific patterns in eDNA distribution, samples along the same transect and at the same sites were taken at 1m depth using a pole sampler (n = 5). In addition, deep water samples (2m from the sediment surface) were taken at sampling stations 3 and 4 on the transect using a van Dorn bottle, (n = 2). We also took 2 samples from the shoreline of each lake, and 2 samples from major inflows and outflows that could be identified on each lake (Extended data figure 5). Filtering of water samples was completed within six hours of collection onto 47mm GF/F filters (ThermoFisher Scientific; nominal pore size = 0.7µm). Filters were dry frozen at -20°C and transported to McGill University on dry ice for molecular analysis.

### eDNA molecular analysis

DNA was extracted from filters using the Qiagen Blood and Tissue kit with some modifications to the manufacturer’s instructions (Supplementary Note 2). Extractions were treated with the OneStep PCR Inhibitor Removal Kit (Zymo Research, Irvine, California).

We created amplicon libraries with two markers; the COI marker was used for targeting broader eukaryotic biodiversity^41^ and the 12S marker was used to characterise the fish assemblages^42^. DNA was amplified in triplicate 12.5µl reactions with some changes from the original publications (Supplementary Note 2). PCR amplicons for each sample were combined, cleaned with AMPure beads and indexed with the Nextera DNA indexing kit for 96 samples (Illumina). A second clean-up with AMPure beads was performed, and libraries were quantified and normalised to 5ng/ul. A mock community of North American fish species was sequenced alongside our samples to evaluate the efficiency of our molecular methods and bioinformatics steps (Extended data table 3). Equimolar amounts of DNA were combined and a total of 433 samples were allocated across five sequencing lanes and sequenced with even depth per sample. Sequencing was conducted using 2×300bp Illumina MiSeq at the McGill University and Génome Québec Innovation Centre, Montréal.

### Bioinformatics

Génome Québec, Montréal, provided a set of demultiplexed sequences that were used for the bioinformatics analyses. We used a denoising pipeline to filter errors and cluster sequences into amplicon sequence variants or ASVs^43^. This approach includes sorting the sequences into markers (12S and COI sequences), adapter removal, quality filtering, merging and quality control (Supplementary Note 3). We filtered the merged and vetted sequences, based on a sequence length ± 20bp around the target amplicon size (152-192bp for 12S, 293-333bp for COI; Supplementary Note 3). The remaining sequences were dereplicated, low-abundance sequences were removed and ASVs were created (Supplementary Note 3). Finally, all reads were aggregated into a count matrix for analysis that gives the number of reads per sample per ASV. This pipeline can be found at https://github.com/CristescuLab/YAAP.

We assigned taxonomy to the ASVs using BLAST+^44^ with high stringency parameters (98% identity, 90% query coverage for 12S, 95% identity, 95% query coverage for COI) and used the last common ancestor algorithm in BASTA^45^ to assign taxonomic identity (Supplementary Note 3). For the 12S marker, we matched to a local database composed of sequences from fish species during the last 50 years of monitoring data at the IISD-ELA as well as government surveys of the same region. For the COI marker, we used a local copy of the NCBI database (downloaded on the 12 August 2018). We then created a subset of zooplankton taxa that had been assigned at least family-level taxonomic identity in the following groups: Amphipoda, Calanoida, Cyclopoida, Diplostraca, Chaoboridae, Rotifera, Ostracoda and Trichoptera. We performed an adjustment of sequence numbers based on the small amount of contamination that appeared in the mock community; this was done by calculating proportions for each species that would remove the contamination from the mock community and the negative controls and then adjusted this to every sample’s library size. We then subtracted this amount of sequences from every library.

### Statistics

We used the vegan v 2.5-2^46^ package in R (v 4.0.2) to compute diversity statistics and visualise species accumulation curves. We performed the following analyses to address our original objectives:

### 1. Within-lake variation in eDNA distribution

To examine the contributions to within-lake patchiness in eDNA distribution, we initially analysed whether eDNA sample location in the lake (e.g. shoreline, deep-water, pelagic-surface transect, in- and out-flows) influenced the recovered eDNA community composition by performing PERMANOVA using a Bray-Curtis dissimilarity matrix on the ASV abundance x sample table with sample location as the explanatory variable. Separate models were created for fish and zooplankton data. Samples were permuted 999 times with lake identity as a strata effect.

We then conducted hypothesis testing on per-sample ASV abundances in relation to typical habitat use by each species. Fish and zooplankton specialists classified species or higher taxonomic groups according to their habitat use (Fish: littoral-benthic, mid-water benthic, profundal cold water, pelagic; Zooplankton: littoral, profundal cold water, and pelagic (Extended data table 4). We fitted generalised mixed effects models using glmmTMB with a zero-inflated negative binomial distribution^47^ to assess the interaction between species habitat use and eDNA sample location (epilimnion, deeper water, shoreline) on per sample ASV abundances for each species. Separate models were created for fish and zooplankton. We accounted for effects that might be due to differing library sizes by including this as an offset term. We allowed intercepts to vary according to lake and species by including these as crossed random effects. We assessed the importance of the interaction between habitat preference and sample location in predicting the ASV abundances by testing significance using a likelihood ratio test with a Chi-squared distribution. We used the DHARMa v0.4.4 package to test for overdispersion, correct handling of zero-inflated data, and model assumptions^48^. Finally, we visually explored the contribution of lake size and lake state (i.e. stratified or mixed) to the distribution of eDNA at different sample locations within the lakes.

### 2. Between-lake variation in species detection

We investigated our model predictions (Figure 1A), which describe the role of freshwater connectivity in explaining expected and unexpected species detections made with eDNA. We categorised fish and zooplankton eDNA detections as unexpected if they were not predicted by population records from current and historical surveys for the lake in question. We created statistical models with the number of species detections per sample as the response variable. Each water sample provided two datapoints, one count of expected detections and one count of unexpected detections. We therefore included the filter identity as a random effect. We included sample location (i.e., pelagic, deep water, shoreline), lake chain number and count type (i.e. expected or unexpected detections) as explanatory variables, as well as the two-way interactions between sample location and count type, and lake chain number and count type. Including these two-way interactions would investigate whether certain sample locations are predisposed to give more unexpected detections. It would also investigate whether increasing connectivity (i.e. lakes downstream in the lake network) would increase unexpected eDNA detections.

Initially we also included the number of inflows to a lake in the model, but we found that this explained the same proportion of variation as lake chain number and therefore removed this term. Because species detection is likely to increase with increasing library size, we also included scaled library size as a covariate in the model. We fitted two series of models in glmmTMB using the negative binomial family (for the fish dataset) and the poisson family (for the zooplankton dataset). We tested for overdispersion and model assumptions using the DHARMa v0.4.4 package to confirm that these were the best distributions to use with the respective datasets^48^. We confirmed the significance of the fixed effects terms using a likelihood ratio test with a Chi-squared. Model reduction was performed to remove any non-significant terms, although in all cases we retained library size as a covariate, because this is a requisite part of our experimental design. We used the emmeans v1.7.0 package to perform post-hoc tests to investigate the differences in how detections accumulated in lake networks for expected and unexpected detections.

### 3. Stream discharge and eDNA detections

We investigated the role of stream discharge in explaining eDNA detections in inflows that did not match detections from conventional methods. We hypothesised that streams with a greater discharge would transport more eDNA from upstream lakes, resulting in greater numbers of unexpected detections in receiving lakes. We created a subset of the dataframe with expected and unexpected detections that only included samples from these stream inflows. Discharge was measured either using weirs between lakes or by placing a current flow meter (Gurley Precision Instruments, Troy NY) at five points across the width of each stream and calculating discharge in cms^49^. We then created negative binomial glmmTMB models for fish and zooplankton datasets as before, investigating the interaction between discharge and count type (i.e. expected and unexpected) on the number of detections in the samples. As before, we confirmed the significance of the fixed effects terms using a likelihood ratio test with a Chi-squared distribution and used the DHARMa package to test for model assumptions^48^. We retained library size as a covariate in all models.

