## Extended data figures and tables for "Freshwater connectivity transforms spatially integrated signals of biodiversity"

Extended data file

Joanne E. Littlefair*, José S. Hleap, Vince Palace, Michael D. Rennie, Michael J. Paterson, Melania E. Cristescu
*corresponding author

Table of contents


| Item | Page number |
|---|---|
| Extended data table 1: Sequences retained at each stage of bioinformatics analysis | 2 |
| Extended data table 2: Physical properties of lakes and lake networks | 3 |
| Extended data table 3: Mock community of fish tissues | 4 |
| Extended data table 4: Classification of fish and zooplankton ASVs according to species habitat use | 6 |
| Extended data figure 1: Sample accumulation curves for 12S and COI ASVs | 8 |
| Extended data figure 2: Class level classification of ASVs for A) 12S and B) COI markers | 9 |
| Extended data figure 3: location of fish ASVs according to species habitat preferences and lake size | 10 |
| Extended data figure 4: location of zooplankton ASVs according to species habitat preferences and lake size | 11 |
| Extended data figure 5: Sampling schematic | 12 |

Extended data table 1: Sequences retained at each stage of bioinformatics analysis

Table shows the sequences retained at each stage of bioinformatics analysis. We received a total of 85,504,066 raw reads (201,186 average per sample) from Génome Québec in the form of demultiplexed fastq files. Our pipeline created 585 12S ASVs, of which 154 could be assigned taxonomy at class level and 8,578 COI ASVs, of which 1,356 could be assigned taxonomy at class level (Extended data figure 2). We were able to map the majority of processed sequences back onto these ASVs (95.8% overall for 12S, 88.6% for COI). The 12S marker primarily detected fish: 87.1% of processed 12S sequence counts mapped onto ASVs that were assigned to fish present in the Lake of the Woods area, Ontario.

| | 12S total | 12S mean per sample | COI total | COI mean per sample |
|---|---|---|---|---|
| Raw reads | 85,504,066 | | 85,504,066 | |
| Sequencing adapter trim | 35,204,230 | 82,833 | 18,082,889 | 42,548 |
| After merging | 35,131,597 | 82,663 | 18,037,719 | 42,442 |
| Primer trim | 35,129,369 | 82,657 | 18,035,707 | 42,437 |
| Length filter | 8,443,301 | 19,867 | 18,080,153 | 42,542 |
| Dereplication | 1,122,223 | | 7,120,677 | |
| ASVs created | 585 | | 8,578 | |
| Reads mapped onto ASVs | 8,092,902 (95.8%) | | 16,015,173 (88.6%) | |
| ASVs with taxonomy assigned at class level | 156 | | 1,356 | |
| ASVs with taxonomy assigned at species level | 106 | | 588 | |

Extended data table 2: Physical properties of lakes

Table displays the physical properties of lakes. The depth data refers to the depth measured at sampling point 3 (Extended data figure 5) which was taken at approximately the deepest point of the lake according to historical estimates. The lake size category refers to "s" = small, non-stratified, "mns" = medium, non-stratified, "ss" = small, stratified, "l" = large, stratified. These categories are used on Extended data figures 3 and 4 to classify lakes.

| Network | Lake ID | Measured depth at centre point (m) | Size (ha) | Number of inflows | Position in lake chain | Lake size category* |
|---|---|---|---|---|---|---|
| Chain 1 | 303 | 2.3 | 10 | 0 | 1 | s |
| Chain 1 | 304 | 6.8 | 3 | 0 | 1 | ss |
| Chain 1 | 661 | <1 | NA | 1 | 2 | s |
| Chain 1 | 470 | 1.6 | 6 | 1 | 3 | s |
| Chain 1 | 239[a] | 32.8 | 54 | 3 | 1 | l |
| Chain 1 | 240[a] | 14 | 44 | 2 | 4 | ss |
| Chain 1 | 979[b] | 0.9 | 2 | 1 | 5 | s |
| Chain 1 | 663 | 5.9 | 47 | 2 | 6 | mns |
| Chain 1 | 664 | 13.5 | 67 | 2 | 7 | l |
| Chain 2 | 114[a] | 2.45 | 12 | 0 | 1 | s |
| Chain 2 | 115 | 1.3 | 77 | 1 | 1 | s |
| Chain 2 | 467 | 13.9 | 210 | 4 | 2 | l |
| Chain 2 | 665[b] | 11.5 | 30.3 | 2 | 3 | ss |
| Chain 2 | 164 | 6.7 | 20 | 2 | 5 | ss |
| Chain 2 | 165 | 4.4 | 18 | 2 | 4 | mns |
| Chain 3 | 429[b] | 7.5 | 16 | 0 | 1 | mns |
| Chain 3 | 628[b] | 17.5 | 19.4 | 1 | 2 | ss |
| Chain 3 | 626[a] | 11.9 | 25.9 | 0 | 1 | ss |
| Chain 3 | 627[b] | 8.2 | 35.5 | 1 | 3 | mns |
| Chain 3 | 625[b] | 35.4 | 78.3 | 5 | 4 | l |
| Chain 3 | 653 | 26.2 | 182.9 | 4 | 5 | l |

[a]Lakes with regular/annual monitoring for at least 10 years

[b]Sampled for fish between 2014-2018 using modified broad scale monitoring method

Extended data table 3: Mock community of fish tissues

We evaluated the ability of several primer pairs against tissue extracts of 40 North American fish species obtained from the Ministère des Forêts, de la Faune et des Parcs (Québec). The selected species were chosen to represent a diversity of families within the Actinopterygii, to assess how our taxonomic assignment method performs when only congeneric reference sequences were available in the reference data base, and to include some species known to be present at the IISD Experimental Lakes Area. Muscle or fin tissue was extracted using Qiagen Blood and Tissue kits and equimolarised to 15ng/µl. Mock community samples were included in the sequencing to evaluate the efficiency of our molecular methods and bioinformatics steps. Species were identical to those used in primer testing with the addition of one fish (*Etheostoma nigrum*) for this sequencing step only, for a total of 41 species. Equimolar amounts of DNA were combined and two replicate mock community libraries were PCR amplified and sequenced. We detected 33 of the 41 fish species in the mock community by metabarcoding the 12S fragment. Of the eight species that were not detected, five did not have complete reference sequences in the database

| Species name | Family | 12S reference sequence available | Mock community rep 1 ASV abundance | Mock community rep 2 ASV abundance |
|---|---|---|---|---|
| *Acipenser fulvescens* | Acipenseridae | Yes | 1269 | 772 |
| *Alosa aestivalis* | Clupeidae | Yes | 735 | 389 |
| *Alosa pseudoharengus* | Clupeidae | Yes | 5397 | 3006 |
| *Ambloplites rupestris* | Centrarchidae | Yes | 2823 | 1606 |
| *Ameiurus nebulosus* | Ictaluridae | Yes | 1460 | 841 |
| *Anguilla rostrata* | Anguillidae | Yes | 1057 | 565 |
| *Catostomus commersonii* | Catostomidae | Yes | 2776 | 1555 |
| *Coregonus artedi* | Salmonidae | Yes | 1567 | 860 |
| *Culaea inconstans* | Gasterosteidae | Yes | 7356 | 4331 |
| *Cyprinella spiloptera* | Cyprinidae | Yes | 8764 | 5451 |
| *Cyprinus carpio* | Cyprinidae | Yes | 3454 | 2074 |
| *Dorosoma cepedianum* | Clupeidae | Yes | 1357 | 768 |
| *Esox lucius* | Esocidae | Yes | 607 | 346 |
| *Esox masquinongy* | Esocidae | Yes | 1918 | 1086 |
| *Etheostoma nigrum* | Percidae | Yes | 2766 | 1630 |

| | | | | |
|---|---|---|---|---|
| *Etheostoma flabellare* | Percidae | No – congenerics available | Not detected | Not detected |
| *Fundulus diaphanus* | Fundulidae | Yes | 2632 | 1450 |
| *Hiodon tergisus* | Hiodontidae | Yes | 11983 | 6904 |
| *Labidesthes sicculus* | Atherinopsidae | No | Not detected | Not detected |
| *Lota lota* | Lotidae | Yes | 6943 | 4084 |
| *Luxilus cornutus* | Cyprinidae | Yes | Not detected | Not detected |
| *Microgadus tomcod* | Gadidae | Yes | 4458 | 2655 |
| *Micropterus dolomieu* | Centrarchidae | Yes | 22 | 14 |
| *Morone americana* | Moronidae | Yes | 2691 | 1534 |
| *Morone saxatilis* | Moronidae | Yes | 1085 | 600 |
| *Neogobius melanostomus* | Gobiidae | Unverified mitochondrial sequence | Not detected | Not detected |
| *Notemigonus crysoleucas* | Cyprinidae | Yes | 271 | 155 |
| *Notropis atherinoides* | Cyprinidae | Yes | 4742 | 2936 |
| *Notropis heterodon* | Cyprinidae | Yes | 1243 | 807 |
| *Notropis heterolepis* | Cyprinidae | Yes | 4606 | 2989 |
| *Notropis hudsonius* | Cyprinidae | Yes | 3585 | 2241 |
| *Notropis rubellus* | Cyprinidae | No | Not detected | Not detected |
| *Notropis volucellus* | Cyprinidae | Yes | Not detected | Not detected |
| *Noturus flavus* | Ictaluridae | No | Not detected | Not detected |
| *Osmerus mordax* | Osmeridae | Yes | 800 | 494 |
| *Perca flavescens* | Percidae | Yes | 5042 | 2889 |
| *Percina caprodes* | Percidae | Yes | *Percina macrolepida* detected | *Percina macrolepida* detected |
| *Percopsis omiscomaycus* | Percopsidae | Yes | 12 | 9 |
| *Pimephales promelas* | Cyprinidae | Yes | 3966 | 2460 |
| *Salmo trutta* | Salmonidae | Yes | 1673 | 971 |
| *Sander vitreus* | Percidae | Yes | 12586 | 7519 |

Extended data table 4: Classification of fish and zooplankton ASVs according to species habitat use

Table shows species classified according to habitat preferences; these classifications are used in the within-lake analysis presented in Figure 2. Fish habitat classifications were made based on summer distributions as per Scott and Crossman 1998[†]. Zooplankton habitat classifications were made based on analysis of the long-term monitoring datasets at IISD-ELA.

| Fish species | Habitat classification | Zooplankton taxa | Habitat classification |
|---|---|---|---|
| *Catostomus commersoni* | Littoral-benthic | *Acantholeberis curvirostris* | Littoral |
| *Chrosomus neogaeus/Chrosomus* mitochondrial hybrids | Littoral-benthic | *Aloninae* | Littoral |
| *Coregonus artedi* | Pelagic | *Bosmina/ Bosmina longirostris* | Pelagic |
| *Coregonus clupeaformis* | Profundal | *Ceriodaphnia* | Littoral |
| *Cottus cognatus* | Profundal cold water | *Chaoborus punctipennis Chaoborus/Chaoboridae* | Pelagic |
| *Couesius plumbeus* | Pelagic | *Chydorus/Chydoridae* | Littoral |
| *Culaea inconstans* | Littoral-benthic | *Cypria* | Littoral |
| *Esox lucius* | Littoral-benthic | *Diaphanosoma birgei* | Pelagic |
| *Etheostoma exile* | Midwater benthic | *Diporeia hoyi* | Profundal cold water |
| *Etheostoma nigrum* | Littoral-benthic | *Epischura lacustris* | Pelagic |
| *Lota lota* | Profundal cold water | *Euchlanis dilatata* | Littoral |
| *Margariscus natchtriebi* | Littoral-benthic | *Eurycercus longirostris* | Littoral |
| *Notropis atherinoides* | Pelagic | *Holopedium glacialis* | Pelagic |
| *Notropis heterolepis* | Littoral-benthic | *Kellicottia* | Pelagic |
| *Notropis hudsonius* | Pelagic | *Leptodiaptomus sicilis* | Profundal cold water |
| *Perca flavescens* | Littoral-benthic | *Leptodiaptomus minutus* | Pelagic |
| *Percina caprodes* | Littoral-benthic | *Leptodora kindtii* | Pelagic |
| *Percopsis omiscomaycus* | Littoral-benthic | *Limnocalanus macrurus* | Pelagic |
| *Pimephales promelas* | Littoral-benthic | *Macrothrix* | Littoral |

| | | | |
|---|---|---|---|
| *Salvelinus namaycush* | Profundal cold water | *Mochlonyx* | Pelagic |
| *Sander vitreus* | Midwater benthic | *Ophryoxus gracilis* | Littoral |
| *Rhinichthys cataractae* | Midwater benthic | *Pleuroxus* | Littoral |
| | | *Ploesoma hudsoni* | Pelagic |
| | | *Polyarthra dolichoptera* | Profundal cold water |
| | | *Polyphemus pediculus* | Littoral |
| | | *Scapholeberis* | Littoral |
| | | *Simocephalus* | Littoral |
| | | *Skistodiaptomus oregonensis* | Pelagic |
| | | *Trichocerca capucina* | Pelagic |

† Scott WB, Crossman EJ. 1998. Freshwater fishes of Canada. Galt House Publications. Oakville, ON, Canada.

Extended data figure 1: Sample accumulation curves for 12S and COI ASVs

Lake sample accumulation curves in the three lake networks for two markers: A) Chain 1, 12S marker, B) Chain 1, COI marker, C) Chain 2, 12S marker, D) Chain 2, COI marker, E) Chain 3, 12S marker, F) Chain 3, COI marker. See extended data table 4 for the classifications of the lakes within each of the three networks. Each lake at the IISD Experimental Lakes Area has a unique identification number and these are represented in the legends.

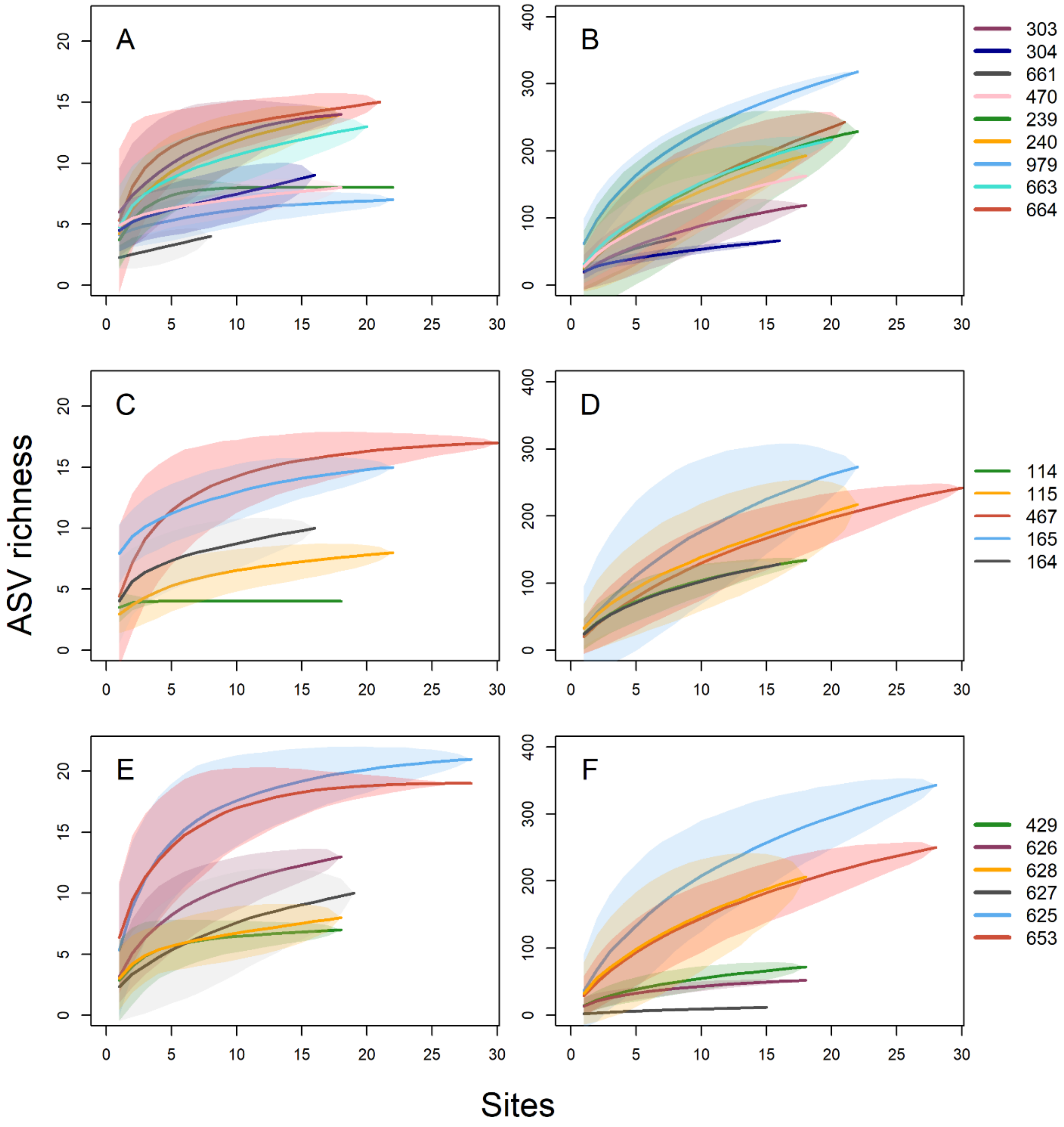

Extended data figure 2: Class level classification of ASVs for A) 12S and B) COI markers

Pie charts show incidence of class level assignments of ASVs when blasted against a universal nucleotide database downloaded from NCBI. Numbers represent counts of taxa assigned within that class. Even though the 12S marker was specifically designed for fish, it detected a variety of other taxa.

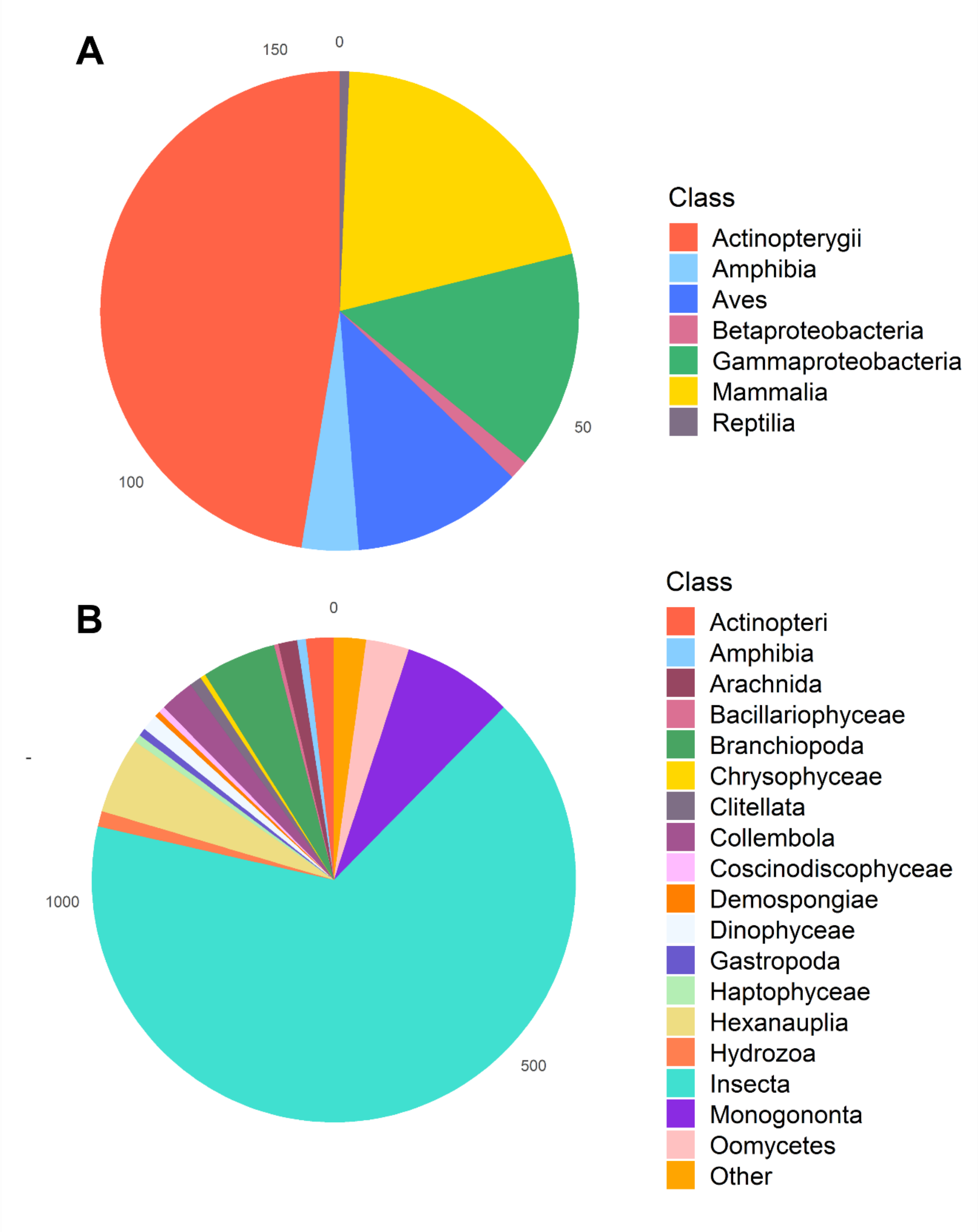

Extended data figure 3: location of fish ASVs according to species habitat preferences and lake size

Bars show the average proportion of fish ASVs per water sample weighted by abundance in different sized lakes. The x axis shows the position of the water samples within the lakes. Different facets display lakes of differing sizes and states. Small, non-stratified lakes were defined as being ≤12ha. Medium non-stratified lakes were 16-47ha. Small, stratified lakes were defined as 3-44ha and were usually deeper than the small lakes in order to stratify. Large, stratified lakes were ≥54ha. There were approximately even numbers of lakes in each group (see Extended data table 2 for the categorisation of lakes into these groups). Bar graph segments are coloured in themes according to the species habitat preferences displayed in Figure 2, i.e. blue = pelagic species, yellow = littoral species, red = profundal species, and green = midwater-benthic species.

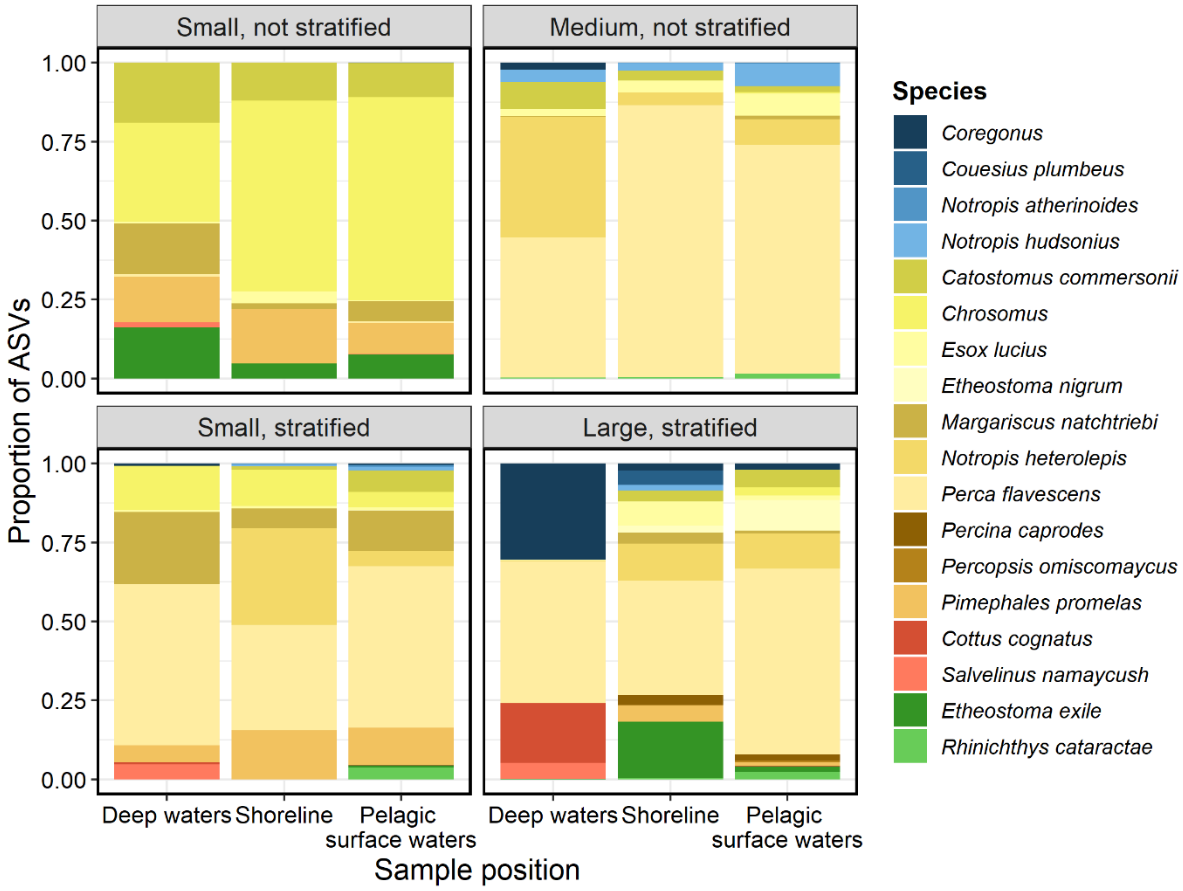

Extended data figure 4: location of zooplankton ASVs according to species habitat preferences and lake size

Bars show the average proportion of fish ASVs per water sample weighted by abundance in different sized lakes. The x axis shows the position of the water samples within the lakes. Different facets display lakes of differing sizes and states. Small, non-stratified lakes were defined as being ≤12ha. Medium non-stratified lakes were 16-47ha. Small, stratified lakes were defined as 3-44ha and were usually deeper than the small lakes in order to stratify. Large, stratified lakes were ≥54ha. There were approximately even numbers of lakes in each group (see Extended data table 2 for the categorisation of lakes into these groups). Bar graph segments are coloured in themes according to the species habitat preferences displayed in Figure 2, i.e. blue = pelagic species, yellow = littoral species, and red = profundal species.

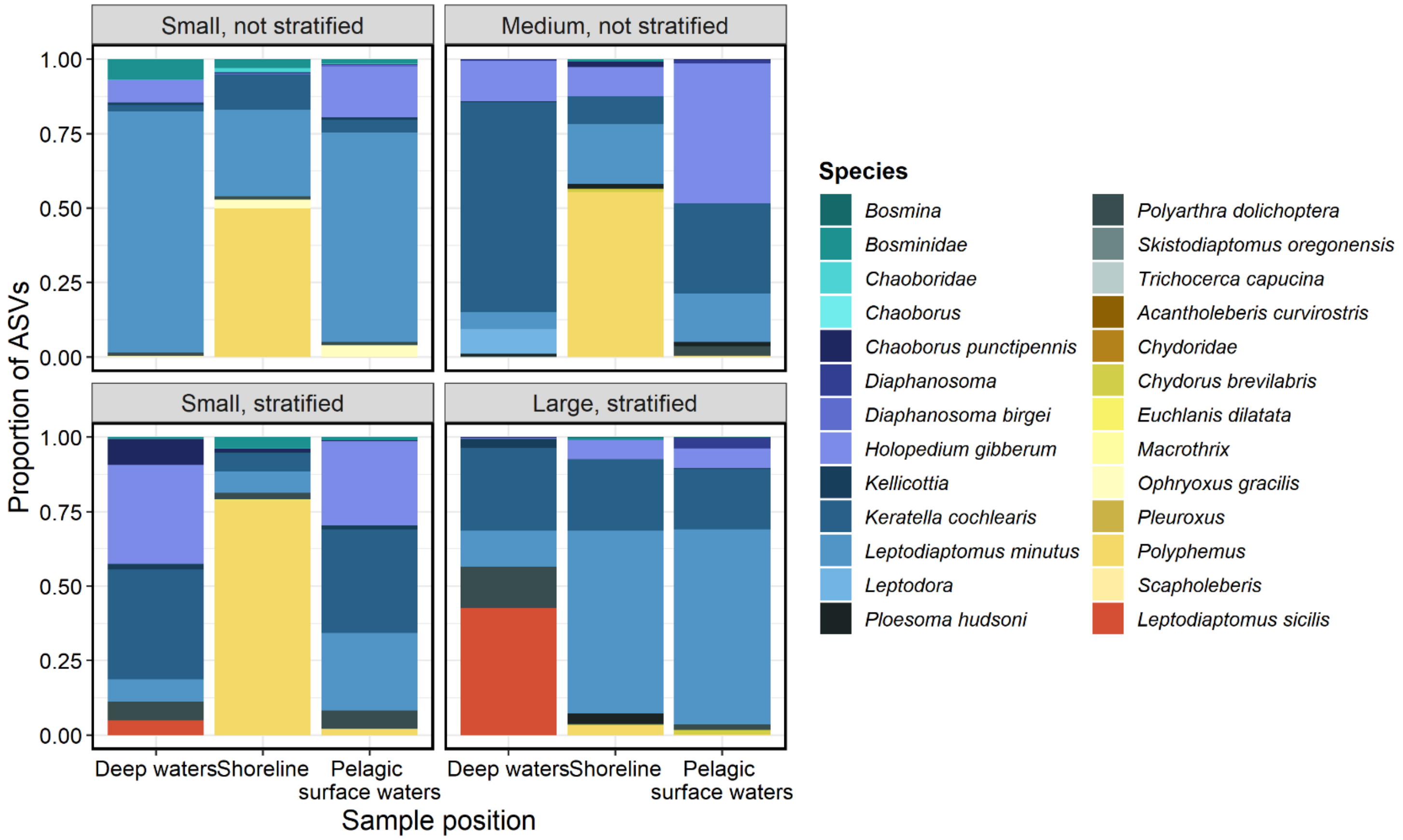

Extended data figure 5: Sampling schematic

Sampling strategies were designed to maximise the coverage of different habitats within the lakes. Circles on the diagram represent where water samples were collected. A transect of five evenly spaced points was designed across the width of each lake, with the third point situated at the deepest point in the lake. On the transect, samples were taken at the surface and 1m below the surface using a pole sampler. At the third point a zooplankton haul was also collected. Two samples were also taken in the deep water of each lake using a van Dorn bottle, two metres from the lake bed, at transect points three and four. Not all lakes were stratified, but we collected samples from the depths in all lakes. Two samples were also taken at the shoreline of each lake, away from in- and outflows. Two samples were taken at each in- and outflow.

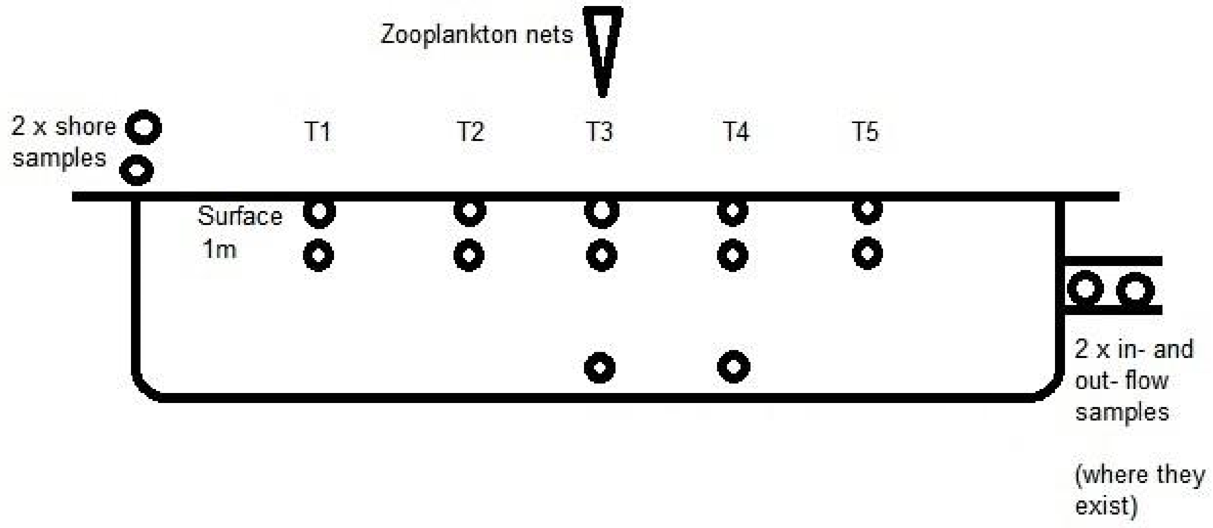
