## Supplementary Information for "Freshwater connectivity transforms spatially integrated signals of biodiversity"

Joanne E. Littlefair*, José S. Hleap, Vince Palace, Michael D. Rennie, Michael J. Paterson, Melania E. Cristescu

*corresponding author

Table of contents


| Item | Page number |
|---|---|
| Supplementary Note 1: Broad scale monitoring of fish and zooplankton populations | 2 |
| Supplementary Note 2: eDNA molecular work and strategies for minimising contamination | 4 |
| Supplementary Note 3: Taxonomic assignment and processing of sequence data | 6 |
| Additional references | 8 |

Supplementary Note 1: Broad scale monitoring of fish and zooplankton populations

Lakes were sampled for fish community composition using several different methods. Lakes that were monitored annually to bi-annually for fish (Extended data table 2) used a combination of non-lethal gillnetting and trapnetting. During spring and autumn, Beamish-style trap nets[1] were deployed with either a central lead set perpendicular to shore, or without a central lead with one wing tied to shore and set roughly parallel to the shoreline. Holding pots of both nets were suspended by floats and the pot and wings were secured with anchors. Holding pots ranged in volume from 2.2 to 6.1 $m^2$ and were typically set in 2–4 m of water. Trap nets were set in fixed locations for 4–6 weeks and emptied every 2–5 days. Mesh sizes of trap nets ranged from 3 to 6 mm, and the smallest fish retained ranged from 19 to 35 mm. In the autumn, short set (approximately 20 minutes) gillnets were deployed on lake trout spawning shoals. All fish from either method are returned to the lake alive after weighing, measuring and tagging.

Since 2014, several lakes have been sampled using a modified version of the Ontario Ministry of Natural Resources Broadscale Monitoring method (Extended data table 2), which applies both North American (25 m, 38–127 mm) and Ontario small mesh (12.5m, 13–38 mm) nets and are area-weighted by depth strata[2]. The density of nets within a depth stratum are dictated by the area of the strata represented in the lake.

All other lakes have been surveyed 1–2 times using a set of experimental gillnets, beach seines, trap nets and minnow traps. Surveys typically lasted 2–4 days and involved intensive sampling of the ecosystem. Between 1–6 panels of 50 m length gillnets ranging in mesh size from 19 to 130 mm were set overnight and retrieved the following day[3]. Trap nets (of the style described above) and minnow traps were typically fished for 1–2 days.

Historical monitoring information on zooplankton at the IISD-ELA was collected using 3 main techniques. Zooplankton in lakes 114, 164, 165, 303, 304, and 979 were collected using a flexible 7cm diameter flexible hose deployed from the water surface to just above the sediments at multiple stations in each lake[4–6]. Samples from different stations were combined and sieved with a 53 µm net before counting. Lakes 239, 240, 625, and 626 were sampled using 2 vertical hauls of a double-barrelled 53 µm net lowered to just above the sediments at the deep station of each lake[7] while lakes 429, 470, and 627 were sampled using 2 vertical hauls of a 30 cm diameter 53 µm Wisconsin net at the deep station of each lake. All samples were preserved in 4% formalin after narcotization with methanol. In each sample, at least 300 crustacea and rotifers were identified to the lowest taxonomic category. These data are in addition to zooplankton hauls taken in 2017 using techniques described in the main manuscript.

Supplementary Note 2: eDNA molecular work and strategies for minimising contamination

Filters were extracted using the Qiagen Blood and Tissue kit. We followed the manufacturer's instructions with some modifications: 370µl buffer ATL was used in the initial incubation step, filters were incubated in ATL and proteinase K for 16 hours overnight, and the DNA was eluted in 2 x 40µl of AE buffer and stored at -80ºC after elution. DNA was stored at -80ºC after elution. DNA was amplified in triplicate 12.5µl reactions using COI[8] and 12S markers[9] selected to target broader biodiversity (COI: 313bp) as well as fish assemblages (12S: 163–185bp). For the 12S marker we used 7.4µl nuclease free water (Qiagen), 1.25µl 10X buffer (Genscript), 1mM $MgCl_2$ (ThermoFisher Scientific), 0.2mM GeneDirex dNTPs, 0.05mg bovine serum albumen (ThermoFisher Scientific), 0.25mM each primer, 1U *taq* (Genscript) and 2µl DNA in a final volume of 12.5µl. For the COI marker, the mastermix contained 7.875µl nuclease free water, 1.25µl 10X buffer (Genscript), 1mM $MgCl_2$ (ThermoFisher Scientific), 0.1mM GeneDirex dNTPs, 0.0125mg bovine serum albumen (ThermoFisher Scientific), 0.2mM each 10mM primer, 1.25U *taq* (GenScript) and 2µl DNA in a final volume of 12.5µl. The thermocycling regime followed that of the original paper. Amplicons were run on 1% agarose gel stained with SYBR™ Safe DNA Gel Stain (Thermo-Fisher Scientific) and visualized with UV light.

We structured field and laboratory work to minimise contamination between samples and lakes, as eDNA transfer between lakes was one of the fundamental research questions of our study. We sampled one lake per day and cleaned all sampling equipment thoroughly each evening. The sampling pole and van Dorn bottle were cleaned with 20% bleach between within-lake sampling points, and thoroughly cleaned with 20% bleach and soapy water at the end of each day. New gloves were used for the collection of every sample. All epilimnetic, deep-water, shoreline and in- and out-flow samples were taken in single-use whirlpak bags, which were double bagged inside a large ziplock bag. Each day, a field negative blank of autoclaved distilled water was transported into the field and filtered back in the laboratory using the same procedure as the field samples. Filtration equipment was scrubbed with hot soapy water and soaked for >10 minutes in 30% bleach, triple rinsed with distilled water, and autoclaved before reuse. Samples were filtered in a room that had never previously been utilised for animal tissue or DNA work. Before work began, all floors were cleaned with floor cleaner, laboratory coats washed, and surfaces and equipment wiped down with 20% bleach. Filters were stored at -20°C in a freezer that was not used for storing animal tissue at the IISD-ELA.

DNA extraction and pre-PCR laboratory work took place in a dedicated eDNA facility at McGill University. For both DNA extraction and library preparation, only one lake was processed per day. All equipment and surfaces were wiped with 20% bleach before each use. These steps minimised potential cross-lake contamination from open tubes and multichannel pipettes. Filter tips were used at all stages of molecular work. Both negative DNA extraction and PCR controls were included for every plate by substituting with nuclease free water (Qiagen). All filtration, extraction, and PCR negative controls were amplified in triplicate.

Almost all field, extraction, and PCR negative controls showed no amplification on gel images. Sequencing of negative controls showed a small number of sequences remained; these were removed during the bioinformatic adjustment to remove contamination.

Supplementary Note 3: Taxonomic assignment and processing of sequence data

Our goals for taxonomic assignment were to examine species detection in the IISD Experimental Lakes Area/Lake of the Woods area with eDNA as well as evaluating the taxonomic breadth of the primers. We used an amplicon sequencing variant (ASV) approach which filtered and denoised sequences to achieve this. Demultiplexing was performed by Génome Québec, Montréal. We used seqkit v1.2[10] to sort 12S and COI sequences and process them separately. Adapters were removed using paired-end trimming with cutadapt v1.18, using a quality threshold of 25 and removing undefined bases at the end of sequences[11]. Reads were merged with pear v0.9.10[12] with a quality score of 20 and discarded if merged reads were less than 100bp. Any remaining gene-specific primers were evaluated and removed. We examined the resulting base quality scores with fastqc v0.11.5[13]. Finally, we filtered by sequence length ± 20bp around the target amplicon size (152-192bp for 12S, 293-333bp for COI). We dereplicated sequences with VSEARCH 2.9.1[14]. We then created ASVs with unoise3[15] from USEARCH v10.0.240[16], discarding all ASVs with a minimum size of less than 8 copies. Finally, all reads were aggregated into a count matrix that gives the number of reads per sample per ASV.

We then performed two BLAST analyses on the 12S ASV sequences. We created a local database of fish species known to exist at the Experimental Lakes Area/Lake of the Woods area from 50 years of broad-scale monitoring data and local government surveys. We assigned taxonomy to the 12S ASVs using BLAST+ v 2.2.26[17] with high stringency parameters (98% identity, 90% query coverage) and used the last common ancestor algorithm in BASTA v1.3.2[18] to assign taxonomic identity. These parameters had been shown by analyses of the mock community to optimise recovery and species discrimination for North American freshwater fish (Extended data table 3). We then grouped together ASVs which matched to the same fish species, as populations might have more than one haplotype (no ASV matched to more than one fish). We performed a second BLAST on the 12S ASVs to evaluate general biodiversity and the taxonomic breadth of the primers, using a local copy of the NCBI nucleotide database downloaded on 12 August 2018, and assigned taxonomy to sequences using the LCA algorithm in BASTA. We assigned taxonomy to the COI marker using the local copy of the NCBI database and assigned taxonomy using BASTA with 95% percent identity and 95% query coverage. We then targeted zooplankton taxa by retaining only ASVs which matched to the following taxonomic groups: Amphipoda, Calanoida, Cyclopoida, Diplostraca, Chaoboridae, Rotifera, and Ostracoda.

*Species detection with eDNA*

For two groups of congenerics we accepted detection at the genus level. For the *Chrosomus* genus, hits matched to *Chrosomus neogaeus*, despite the fact that “cybrids” – fish which have a *C. eos* nuclear genome with a *C. neogaeus* mitochondria also exist at ELA, as well as hybrids between the two species[19]. However, as both of our markers were mitochondrial, we could not distinguish between *C. neogaeus* and the mitochondrial hybrids. Additionally, we detected but could not distinguish between the two fish from the *Coregonus* genus, as reads matched to hits from both species. Our targets, *C. artedi* and *C. clupeaformis,* are known to be closely related and are hybridizing in the Great Lakes[20]. For the purposes of assessing the distribution of eDNA, we recorded detection at the genus level for these two pairs of species. We could detect and differentiate between all other fish species.

*Statistical analysis of ASV abundance in relation to conventional species records*

We analysed the number of per-sample, per-species ASV counts in lakes where conventional monitoring records confirmed the presence of species. We used a quasipoisson model to account for overdispersion with ASV count number as the response variable and conventional monitoring records as a binary presence/absence predictor.
